# Extracellular matrix mechanical cues (dys)regulate metabolic redox homeostasis due to impaired autophagic flux

**DOI:** 10.1101/2025.01.09.631485

**Authors:** Heloísa Gerardo, Tânia Lourenço, Júlio Torres, Manuela Ferreira, Célia Aveleira, Susana Simões, Lino Ferreira, Cláudia Cavadas, Paulo J. Oliveira, José Teixeira, Mário Grãos

## Abstract

Extracellular matrix (ECM) stiffness is increasingly recognized as a critical regulator of cellular behavior, governing processes such as proliferation, differentiation, and metabolism. Neurodegenerative diseases are characterized by mitochondrial dysfunction, oxidative stress, impaired autophagy, and progressive softening of the brain tissue, yet research into how mechanical cues influence cellular metabolism in this context remains scarce. In this study, we evaluated the long-term effects of brain-compliant, soft ECM on mitochondrial bioenergetics, redox balance, and autophagic capacity in neuronal cells. Using human neuroblastoma (SH-SY5Y) and mouse hippocampal (HT22) cell lines, as well as primary mouse neurons, we observed that prolonged exposure to soft ECM resulted in mitochondrial bioenergetic dysfunction, redox imbalance, and disrupted autophagic flux. These findings were consistently validated across both human and mouse neuronal cells. Our data indicate a decreased maximal autophagic capacity in cells exposed to long-term soft ECM, potentially due to an imbalance in autophagosome formation and degradation, as demonstrated by decreased LC3 II levels following chloroquine-induced autophagic flux inhibition. This impairment in autophagy was coupled with increased cellular oxidative stress, further indicating metabolic alterations. These findings emphasize the critical role of ECM stiffness in regulating neuronal cell metabolism and suggest that prolonged exposure to soft ECM may mimic key aspects of neurodegenerative disease pathology, thereby enhancing the physiological relevance of *in vitro* models. This study underscores the necessity for further research into ECM mechanics as a contributing factor in neurodegenerative disease progression and as a potential target for therapeutic strategies.

## Introduction

Different tissues and organs show a specific composition of their extracellular matrix (ECM), which results in distinctive biomechanical properties. These are essential not only for structural integrity, but also for controlling different aspects of cell biology in physiology and disease [1, 2]. Cells sense ECM stiffness through integrin-based adhesions, which facilitate the generation of intracellular tension via the actomyosin cytoskeleton. This mechanical input is subsequently converted into biochemical signals that regulate various cellular processes, including proliferation, differentiation, metabolism, and gene expression. [2-4]. In the nervous system, mechanotransduction plays a particular critical role, since processes such as axon guidance, stem/progenitor cell differentiation and synapse maintenance are regulated by the mechanical properties of the extracellular environment [1, 5-8]. Therefore, changes in ECM composition and stiffness during development, aging, and neuropathology alter mechanical inputs sensed by neurons and glial cells, impacting the overall functionality of the central nervous system and potentially contributing to the progression of neurodegenerative diseases [5, 6, 9]. Furthermore, neurodegenerative diseases are characterized by mitochondrial dysfunction, oxidative stress, and impaired autophagy [10-13], along with progressive loss of brain tissue stiffness [14-17], making this a suitable pathological context for investigating the interplay between cell mechanics and metabolism.

Recent studies have highlighted substantial alterations in cancer cells’ metabolic processes in response to low actomyosin tension favoring conditions, such as soft ECM [4, 18, 19]. Key findings include increased mitochondrial fission and reactive oxygen species production [20-22], neutral lipid and cholesterol synthesis [23, 24], as well as impaired autophagosome formation [25, 26]. Although the interplay between mechanotransduction and metabolic regulation has been deeply investigated in cancer, those mechanisms in neural cells, often used *in vitro* to mimic neurodegenerative conditions, remain poorly understood.

Using CG-4 cells and rat oligodendrocyte precursor cells (OPCs), we previously demonstrated that brain-like ECM stiffness enhances cell differentiation and maturation into oligodendrocytes, a critical cell type impacted in multiple sclerosis. These findings suggest that brain-compliant ECM optimally supports these processes, underscoring the importance of replicating *in vivo* environments [27]. Additionally, human Wharton’s jelly mesenchymal stem cells (WJ-MSCs) cultured on softer ECM showed increased euchromatin and altered gene expression, further highlighting the influence of ECM stiffness on cellular function [28]. Building on these findings, we investigated whether prolonged ECM mechanical cues influence mitochondrial bioenergetics, redox balance, and autophagy in neuronal cells. Our results show that long-term exposure to brain-compliant soft ECM (1-2.5 kPa), which has significantly lower stiffness compared to standard culture conditions (∼GPa), affects mitochondrial bioenergetic (membrane polarization and oxygen consumption), resulting in reduced metabolic activity and energy production in neuronal cells. These alterations were associated with redox imbalance and compromised autophagic capacity. Our results further show that the prolonged exposure to soft ECM conditions results in impaired autophagy in neuronal cells, thereby reducing the efficient recycling of subcellular organelles. This disruption leads to mitochondrial bioenergetic dysfunction and redox imbalance. Overall, our study highlights the importance of incorporating physiologically relevant ECM stiffness into studies of neuronal cell metabolism to provide a more accurate *in vitro* model, capturing the cellular, biochemical, and mechanical features of pathological environments observed in neurodegenerative diseases.

## Materials and Methods

### Preparation of polyacrylamide hydrogels (PAHs) and glass coverslips

PAHs [27] were prepared on top of reactive glass coverslips [29] (12 mm or 32 mm diameter for immunocytochemistry and western blot (WB), respectively) [30]. Coverslips were treated with 3-(trimethoxysilyl)propyl methacrylate (Sigma-Aldrich, M6514) diluted 1:200 (v/v) in absolute ethanol, supplemented with 3% (v/v) of diluted acetic acid (diluted 1:10 in water) in order to be chemically activated. Coverslips were rinsed in this solution and allowed to react for 3 minutes, and then rinsed with ethanol and air-dried. For the PAHs production, solutions were prepared by mixing sterile deionized water, 40% acrylamide (Bio-Rad, 161-0140), 2% bis-acrylamide (AppliChem, A3657), and 0.01% (v/v) TEMED (Merck, 1,10732,0100) according to **Table 1** to obtain the desired stiffness (elastic modulus - kPa). Before proceeding, the pH of the polyacrylamide solution was adjusted to 7.5 by adding HCl (2 N) and the solution was degassed for 30 minutes [31], using a vacuum system. Next, N-Acryloxysuccinimide ester — NHS (Acros Organics, 10085702) (20 mg/mL) prepared in toluene was added to allow the subsequent functionalization of the hydrogels with proteins. After degassing, 0.06% (v/v) APS and 22% (v/v) of the NHS solution were slowly added to the acrylamide-containing solution and mixed carefully, to avoid the formation of air bubbles. The polymerization of PAHs was made using an electrophoresis gel casting system with 1 mm spacers (Mini-protean III, Bio-Rad). Both glasses from the casting were chlorosilanated, with 5% (v/v) dichlorodimethylsilane (DCMS) (Sigma-Aldrich) diluted in toluene, for 5 minutes to avoid the adherence of the hydrogels to the surfaces, and then dried with a wipe [32]. The chemically activated coverslips were placed on top of the treated spacer-containing glass (coverslips adhered to the glass surface by using small droplets of water between both glass surfaces), and the system was mounted. The PAH solution was then pipetted carefully between the coverslips adherent to the spacer-containing glass and the outer glass and allowed to polymerize for 30 minutes at RT. The polymerized hydrogels (covalently bound to the coverslips) were removed from the system and washed three times with sterile PBS 1X, 5 minutes each with gentle agitation. Then the PAHs were placed on multiple well plates and sterilized for 30 minutes under UV light. As standard cell culture conditions, glass coverslips (∼GPa) were chemically functionalized, to allow the covalent binding of ECM proteins [28]. Briefly, coverslips were rinsed in 1 M NaOH solution for 30 minutes, then this solution was discarded, and coverslips were incubated with 10% (v/v) (3-Aminopropyl)trimethoxysilane (Alfa Aesar, A11284) in 96% ethanol for 30 minutes. After this incubation, coverslips were washed with abundant water for 3 times, 10 minutes each, with gentle agitation. Then, coverslips were incubated with 3% (v/v) glutaraldehyde (Merck, 1.04239.1000) in PBS 1X for 20 minutes, and washed with abundant water for 3 times, 5 minutes each.

**Table 1.** Composition of hydrogel formulations. Percentage of acrylamide (AC) and bis-acrylamide (BAC) used for the desired elastic modulus (kPa) degrees and volume added (μL) per milliliter of solution.

|  | **1 kPa**<br>(3% AC/0.2% BCA) | **2.5 kPa**<br>(5% AC/0.2% BCA) |
| --- | --- | --- |
| **Acrylamide 40%** | 75 | 125 |
| **Bis-Acrylamide 2%** | 100 | 100 |
| **NHS (20mg/mL)** | 220 | 220 |
| **APS** | 6 | 6 |
| **TEMED** | 1 | 1 |
| **Water** | 598 | 548 |

For selected experiments, commercial easy-coat polyacrylamide hydrogels with a stiffness of 2 kPa (Matrigen) were used.

### Coating

These PAHs and glass coverslips and normal tissue culture polystyrene (TCPs) plates were coated with fibronectin (FN - 5 μg/mL) (Millipore, CC085) and poly-D-lysine (Sigma-Aldrich, P1024) with laminin (Life Technologies, 23017015) (PDL/LN – 10/10 μg/mL), and the proteins were diluted in PBS 1X. PAHs purchased from Matrigen coated with PDL and FN diluted in PBS 1X at a concentration of 5 μg/mL. A combination of PDL/FN was prepared 1:1.

### Neuronal-enriched primary culture

*Ethics:* The animal experimental procedures received approval from the CNC Animal Welfare and Ethics Committee (ORBEA_193_2018/26062018) and the Portuguese Directorate-General for Food and Veterinary (DGAV). All procedures were conducted in strict accordance with the European Union Directive (2010/63/EU) and DGAV general guidelines and were performed by certified personnel. *Primary cell isolation:* Primary cells were isolated from mice brains, males and female pups, between P4-P6 by magnetic activated cell sorting (MACS). Mice pups were euthanized by decapitation and brains were excised and dissociated in single-cell suspension using a Neural dissociation kit (P) (Miltenyi Biotec, 130-092-628) following the manufacturer’s instructions. To remove the meninges, the brains were rolled in tissue paper and rinsed with Hank’s Balanced Salt Solution (HBSS) without calcium and magnesium (Life Technologies, 14185045). They were then dissected into small pieces using a scalpel and transferred to a 15 mL centrifuge tube, with two brains per tube. After a centrifugation of 2 min at 300 g the supernatant was removed and 1950 μL of diluted enzyme P (1900μL of buffer X + 50 μL of enzyme P, pre-heated at 37°C for 15 min) were added and incubated for 15 min in a water bath at 37°C. Then, 30 μL of enzyme A solution (20 μL of buffer Y + 10 μL of enzyme A) was added. The mixture was gently combined, and the tissue was mechanically dissociated by pipetting up and down approximately 10 times using a wide-tipped Pasteur pipette. After that, the mixture was incubated for 10 min in a 37°C water bath, and following incubation, the tissue was gently dissociated by pipetting up and down, using two pipettes with progressively smaller diameters. Incubation in a 37°C water bath for 10 min was made, and then the cell suspension was applied to a 70 μm strainer, pre-washed with HBSS with calcium and magnesium (Life Technologies, 14065049), using 10 times the enzyme volume. The cell suspension was centrifuged at 300 xg for 10 min, then resuspended in HBSS with ions and passed through a 40 μm strainer. Cells were counted. *Immunomagnetic neuronal enrichment:* Cell suspension was centrifuged again at 300 xg for 10 min. Cells were re-suspended in blocking buffer (0.5% (w/v) bovine serum albumin (BSA) in PBS 1X, using 80 μL/ 1x10^6^ cells) and incubated with blocking antibody (10 μL/ 1x10^6^ cells) for 10 min at 4°C. Then, anti-CD140 MicroBeads (Miltenyi Biotec, Germany) (10 μL/ 1x10^6^ cells) was added and incubated for more 15 min incubation at 4°C, to separate the oligodendrocyte precursor cells from the remaining cells. Following this, the cells were washed with blocking buffer and centrifuged at 300 xg for 10 min. MACS LS columns (Miltenyi Biotec, 130-042-401) were placed in the MACS stand and rinsed with 3 mL of 0.5% (w/v) BSA. The cells were resuspended in 0.5% (w/v) BSA (1x10^6^ cells in 500 μL of buffer), loaded onto the column, and subsequently washed 3 times with 3 mL of 0.5% (w/v) BSA. The flowthrough was collected to a centrifuge tube, centrifuged at 300 g for 10 min, and then cells were counted and seeded at a density of 1x10^5^ cells/cm^2^ on glass coverslips and polyacrylamide hydrogels, previously coated with poly-D-lysine and laminin, in Neurobasal medium (Gibco, 21103049) supplemented with supplemented with 10% fetal bovine serum (FBS), (Gibco, 10500064), 1% Penicillin/Streptomycin (Life Technologies, 15140-122), 1% Fungizone antimycotic (Life Technologies, 15290-018), 1% Glutamax (Gibco, 35050038), 20 ng/mL BDNF (PeproTech, 450-02-10uG) and Neurobrew-21 (Miltenyi, 130-093-566).

### Cell culture conditions

Mouse hippocampal neuronal HT22 cell line (obtained from Dr. Dave Schubert from The Salk Institute, La Jolla), human neuroblastoma cell line SHSY-5Y (ATCC, CRL-2266) and primary mouse neurons were used in this study. HT22 cells were cultured in Dulbecco’s modified Eagle’s medium (DMEM) high glucose (Thermo Hyclone, SH30081.01), supplemented with 1% Glutamax (Gibco, 35050038), 10% fetal bovine serum (FBS), (Gibco, 10500064), 1% Penicillin/Streptomycin (Life Technologies, 15140-122) and 1% Fungizone antimycotic (Life Technologies, 15290-018). SHSY-5Y cells were cultured in DMEM-F12 (Gibco, 31331028) medium, supplemented with 10% fetal bovine serum (FBS), (Gibco, 10500064), 1% Penicillin/Streptomycin (Life Technologies, 15140-122) and 1% Fungizone antimycotic (Life Technologies, 15290-018). Primary mouse neurons were cultured in Neurobasal medium (Gibco, 21103049) supplemented with supplemented with 1% Glutamax, 20 ng/mL BDNF (PeproTech, 450-02-10uG), Neurobrew-21 (Miltenyi, 130-093-566), 10% fetal bovine serum (FBS), (Gibco, 10500064), 1% Penicillin/Streptomycin (Life Technologies, 15140-122) and 1% Fungizone antimycotic (Life Technologies, 15290-018). Cells were maintained in a humidified atmosphere (37°C, 5% CO_2_/95% air, 90% humidity).

### Cell culture incubations

HT22 and cells were seeded on PAHs mimicking ECM softening (2 kPa and 1 kPa, respectively) and glass coverslips and allowed to grow for 72 hours. Neuronal-enriched primary cell suspension was directly seeded (1x10^5^ cells/cm^2^) on glass coverslips and PAHs (2.5 kPa) after the cellular isolation. After 3 days in culture (days *in vitro* - DIV), half of the cell culture medium was replaced with fresh medium containing 5-Fluoro-2’-deoxyuridine (5-FDU) (Sigma-Aldrich, F0503) at a final concentration of 10 μM in the well, to inhibit the growth of glial cells. The same procedure was carried out at 5 DIV, and the primary cells – hereafter referred to as primary mouse neurons – were left in culture until 6 DIV (3 before 5-FDU and 3 after). *Mechanotransduction modulation.* Cells on soft substrates were incubated with the RhoA activator CN03 (1 μg/mL, 3 hours) (Cytoskeleton, CN03), which recapitulates the increased cell contractility observed on a stiff substrate. *Autophagic flux measurement.* Cells cultured on PAHs or glass were incubated with the lysosomal protein degradation inhibitor, chloroquine (ChQ; 100 μM, 3 hours) (Sigma-Aldrich, C6628).

### Cellular oxygen consumption measurements

The oxygen consumption of HT22 cells cultured on stiff and soft substrates was assessed using the Resipher device (Lucid Scientific, Atlanta, GA, USA). HT22 cells were seeded (6.2 x 10^3^ cells/well) in pre-coated wells with 70 μL of 2% Matrigel (Corning, 354234) diluted in PBS 1X (stiff) or 30 μl of 100% Matrigel (soft). After allowing the cells to adhere for 1 hour, the plate lid was replaced by the lid with sensors and connected to a central hub computer [33]. After the calibration time, oxygen consumption rate (OCR) was collected in real-time up to 72 hours.

### Mitochondrial area and transmembrane electric potential measurements

HT22 cells were seeded in house-made 2.5 kPa PAHs coated with 5 μg/mL FN (3x10^3^ cells/cm^2^). After 72 hours, cells were incubated with the fluorescent dyes MitoTracker™ Green FM (200 nM, ThermoFisher Scientific, M7514), MitoTracker™ Red FM (100 nM, ThermoFisher Scientific, M7512) and Hoechst 33342 (1 μg/mL, ThermoFisher Scientific, H1399) for mitochondrial network area, membrane polarization, and nuclei, respectively. For semi-quantitative analysis of mitochondrial area and membrane potential (ΔΨm), images were acquired using IN Cell Analyzer 2200 (GE Healthcare) imaging system, using a 40x objective (INCA ASAC 20 x/0.45, ELWD Plan Fluor), with the conjugation of several fluorescence channels according to the different excitation and emission wavelengths.

### Metabolic activity measurements

HT22 cells were seeded in 96-well plate with 2 kPa PAHs PDL/FN coated (5x10^3^ cells/well), and after the 72 hours, the metabolic activity was assessed through resazurin reduction assay [32]. In short, the cell culture medium was removed and replaced by 100 μl of cell culture medium with resazurin (10 ug/ml). The amount of resazurin reduced to resorufin was measured fluorometrically with 570 nm excitation and 600 nm emission wavelengths in Biotek Cytation 3 (Biotek Instruments, Winooski, VT, USA).

### Cellular ATP content measurements

HT22 cells were seeded in a 96-well plate with 2 kPa PAHs PDL/FN coated (5x10^3^ cells/well). After 72 hours, cellular ATP content was measured using CellTiter-Glo Luminescent cell viability assay (G7570, Promega). Briefly, 50 μL of culture medium from each well were replaced by 50 μL of medium coating CellTiter-Glo Reagent. Contents were mixed for 2 minutes to induce cell lysis. Then cell lysate was transferred into a white opaque-bottom 96-well plate and incubated for 10 min at RT. The luminescence signal was obtained in a Cytation 3 multi-mode microplate reader (BioTek Instruments, Inc.).

### Fluorescence-based assessment of intracellular oxidative stress levels

HT22 cells were seeded in 96-well plate with 2 kPa PAHs PDL/FN coated (5x10^3^ cells/well) and in house-made 2.5 kPa PAHs coated with 5 μg/mL FN (3x10^3^ cells/cm^2^). After 72 hours, intracellular ROS levels were assessed using oxidative stress-sensitive report molecule CM-H_2_DCFDA (Life Technologies, C6827). In short, cells were incubated with 5 µM CM-H_2_DCFDA in cell culture medium with 1% FBS and without phenol red, and fluorescence signals were measured with 520 nm excitation and 620 nm emission wavelengths for 1 hour at 37 °C in a Cytation 3 multi-mode microplate reader (BioTek Instruments, Inc.).

### Flow cytometric analysis of intracellular oxidative stress levels

Briefly, cells were incubated with 5 µM CM-H_2_DCFDA, as described above, for 30 min at 37°C. Next, cells were washed with PBS 1X, detached with 0.05% trypsin-EDTA, and resuspended HBSS 1X (with ions). Analysis was performed on a BD Accuri™ C6 (BD Biosciences), ensuring that the entire sample was utilized to maximize the number of cells analyzed, yielding at least 6 x 10^4^ gated events per sample. Cells were gated using SSC-A versus FSC-A to discard cell debris, and on FSC-H versus FSC-A to discard cell doublets. CM-H_2_DCFDA fluorescence was detected using the FL1 channel with a 530/30 nm filter. For each independent biological sample, median fluorescence intensity was measured, with data acquisition based on at least 2.2 x 10^4^ gated events.

### Cellular NAD^+^ and NADH content measurements

HT22 cells were seeded in a 96-well plate with 2 kPa PAHs PDL/FN coated (7.5x10^3^ cells/well). After 72 hours, cellular NAD^+^ and NADH content were measured using the NAD/NADH-Glo™ assay (G9071, Promega) following the manufacturer’s instructions. Briefly, cell culture medium was removed and replaced by 25 μL of PBS 1X and 25 μl of dodecyltrimethylammonium bromide (DTAB). Then a vigorous up and down was made to detach and collect all the cellular content from each well to a respective microtube. A spin down was made in each tube, and their content was separated into two 20 μL containing microtubes, one for NAD^+^ and one for NADH. For the NAD^+^ quantification, 10 μL of HCl (0.4 M) was added and then, both tubes (for NAD^+^ and NADH quantification) were incubated at 60 °C for 15 min. After that, 10 μL of Trizma base was added to the NAD^+^ microtubes, and in microtubes for NADH quantification 20 μL of HCl/Trizma solution were added. Then, all the microtubes were kept at room temperature (RT) for 10 min to cool down. Finally, the content of each tube was pipetted in a white opaque-bottom 96-well plate and the NAD/NADH detection reagent was added to each well in a 1:1 proportion. Subsequently, the luminescence signal was obtained in a Cytation 3 multi-mode microplate reader (BioTek Instruments, Inc.).

### Cellular NADP and NADPH content measurements

HT22 cells were seeded in 96-well plate with 2 kPa PAHs PDL/FN coated, (7.5x10^3^ cells/well). After 72 hours, intracellular NADP^+^ and NADPH levels were measured using the NADP/NADPH-Glo™ assay (G9081, Promega) following the manufacturer’s instructions and as described in the NAD^+^/NADH measurements section.

### Cellular glutathione (GSH) content measurements

HT22 cells were seeded in a 96-well plate with 2 kPa PAHs PDL/FN coated, (7.5x10^3^ cells/well). After 72 hours, cellular GSH content was measured using the GSH/GSSG-Glo™ assay (V6611, Promega) following the manufacturer’s instructions. Briefly, the cell culture medium was removed and replaced by 50 μL of total glutathione reagent. Then, a vigorous shake of 5 min was made to lyse the cells. After that, the cellular content from each well was transferred to a white opaque-bottom 96-well plate. Then 50 μL of luciferin generation reagent was added to each well and after an incubation of 30 min, 100 μL of luciferin detection reagent was added to the mixture. After 15 min incubation the luminescence signal was measured in a Cytation 3 multi-mode microplate reader (BioTek Instruments, Inc.). The GSH levels were determined from comparisons with a linear GSH standard curve, respectively.

### Western blotting analysis

Protein content levels were semi quantitatively analyzed using gel electrophoresis (SDS-PAGE) of whole-cell homogenates, followed by Western blotting with antibodies directed against the denatured form of protein depicted in **Table 2**. To obtain cellular extracts, HT22 (3x10^3^ cells/cm^2^), SHSY-5Y (78x10^3^ cells/cm^2^) cells, and primary neurons (1x10^5^ cells/cm^2^) were seeded in house-made PAHs previously coated with ECM proteins (5 μg/mL FN and 10/10 μg/mL PDL/LN, respectively). After 72 hours, or 6 DIV for primary neurons, cells were collected in Laemmli buffer 2X (120 mM Tris-HCl pH 6.8, 4% Sodium dodecyl sulfate (SDS), 20% glycerol), boiled at 95 °C for 5 minutes for protein denaturation aided by sonication (SONICS Vibracell 750 watt using a cup-horn at 60% intensity for 1 min (10 second on 3 second off cycles). Protein concentration was determined by Pierce 660 nm protein assay (ThermoFisher Scientific). To quantify protein in laemmli buffer, an Ionic Detergent Compatibility Reagent (IDCR) was added to Pierce assay reagent at 50 mg/mL (assay reagent). Then, 10 μL of sample or BSA standards and 150 μL of assay reagent were pipetted in a 96-well plate, incubated for 1 minute with shaking followed by 5 minutes in dark conditions. Afterward, absorbance was read with a wavelength of 660 nm in a Cytation 3 multi-mode microplate reader (BioTek Instruments, Inc.). Protein concentrations were equalized with Laemmli buffer 1X (60 mM Tris-HCl pH 6.8, 5% (v/v) glycerol, 0.6% (v/v) SDS, 0.1M dithiothreitol (DTT) and bromophenol blue) and denatured at 95°C for 5 min. Protein samples (15 µg for HT22 and SHSY-5Y, and 30 µg for mouse primary neurons whole-cell homogenates) were loaded and separated in 7–14% sodium dodecyl sulfate-polyacrylamide gels (SDS-PAGE) followed by transfer to PVDF membranes. Membranes were blocked with 5% fat-free milk or 5% BSA in tris-buffered saline 1X with 0.1% (v/v) tween (TBS-T) for 1 h at RT and incubated overnight at 4°C with primary antibodies. Membranes were further incubated with secondary antibodies (anti-rabbit or anti-mouse) AP-, HRP-, or fluorophore-conjugated (**Table 2**) for 1 h at RT. Membranes were then incubated with enhanced chemifluorescence (ECF) western blotting detection reagent kit (GE healthcare, RPN5785) for AP-conjugated secondary antibodies, or with clarity western ECL substrate (Bio-Rad Laboratories,1705061) for chemiluminescence detection of HRP-conjugated antibodies. The images were acquired using an FX Molecular Imager (Bio-Rad) for chemifluorescence and fluorophore, and a VWR image capture software (V1.8.5.0) for chemiluminescence. The densities of each band were calculated the ImageLab 6.0.1 software (version 2017) from Bio-Rad.

**Table 2.** List of antibodies used for western blotting (WB) and immunocytochemistry (ICC) with references and dilutions.

| Antibody | Dilution | Reference | Company | Identifier (RRID) |
| --- | --- | --- | --- | --- |
| GFAP | 1:1,000 (WB) | Z0334 | Dako | AB_10013382 |
| MAP-2 | 1:1,000 (WB) | sc-74421 | Santa Cruz | AB_1126215 |
| YAP | 1:1,000 (WB)<br>1:200 (ICC) | sc-101199 | Santa Cruz | AB_1131430 |
| PINK1 | 1:1,000 (WB) | ab65232 | Abcam | AB_1603921 |
| Parkin | 1:500 (WB) | 4211 | Cell Signaling | AB_2159920 |
| LC3B | 1:1,000 (WB) | 2775 | Cell Signaling | AB_915950 |
| SQSTM1/P62 | 1:1,000 (WB) | 5114 | Cell Signaling | AB_10624872 |
| ATG7 | 1:1,000 (WB) | 8558 | Cell Signaling | AB_10831194 |
| Beclin | 1:1,000 (WB) | 612113 | BD Biosciences | AB_399484 |
| MTOR | 1:1,000 (WB) | 2972 | Cell Signaling | AB_330978 |
| P-MTOR | 1:1,000 (WB) | 2971 | Cell Signaling | AB_330970 |
| Cofilin-1 | 1:1,000 (WB) | 5175 | Cell Signaling | AB_10622000 |
| P-Cofilin-1 | 1 mg/mL (WB) | 44-1072G | Thermo Fisher | AB_2533565 |
| α-Tubulin | 1:1,500 (WB) | T6199 | Sigma | AB_477583 |
| Anti-rabbit HRP conjugated | 1:5,000(WB) | 7074 | Cell Signaling | AB_2099233 |
| Anti-mouse HRP conjugated | 1:5,000 (WB) | 7076 | Cell Signaling | AB_330924 |
| Anti-rabbit AP conjugated | 1:20,000 (WB) | 111-055-003 | Jackson ImmunoResearch Laboratories, Inc | AB_2337947 |
| Anti-mouse AP conjugated | 1:10,000 (WB) | 115-055-003 | Jackson ImmunoResearch Laboratories, Inc | AB_2338528 |
| Anti-rabbit Cy5 | 1:2,500 (WB) | PA45011 | GE Healthcare | AB_772205 |
| Anti-mouse Alexa 488 | 1:200 (ICC) | A11006 | Invitrogen | AB_2534074 |
| Anti-rabbit Alexa 568 | 1:200 (ICC) | A11036 | Invitrogen | AB_10563566 |

### Immunocytochemistry analysis

After incubation time, cells were fixed using 4% PFA or 4% PFA supplemented with 4% sucrose for primary neurons. Then, the permeabilization and blocking was made using 0.3% PBS-Triton diluted in the blocking solution containing 1% BSA in PBS 1X for 30 min at RT. Next, cells were incubated with primary antibodies (**Table 2**) diluted in blocking solution. Cells were washed three times with PBS 1X and then incubated for 1 hour with appropriate Alexa Fluor-conjugated antibodies (**Table 2**), DAPI (800 ng/mL, D3571, Life technologies).

### Image acquisition and quantitative image analysis

Images were acquired using IN Cell Analyzer 2200 (GE Healthcare) imaging system. For the quantification of fluorescence microscopy images, imaging settings were selected and kept constant during acquisition for all samples. Quantitative image analysis was performed with the ImageJ Fiji (Scion Corporation, USA, RRID:SCR_002285). The YAP nuclear to cytoplasm intensity ratio was automatically calculated using the Intensity Ratio Nuclei Cytoplasm Tool for ImageJ (Intensity Ratio Nuclei Cytoplasm Tool, RRID:SCR_018573).

### Statistical analysis

Statistical analysis was performed using GraphPad Prism version 9.0.0 (GraphPad Software Inc., San Diego, CA, USA). Data were expressed as the mean value ± SEM. The normality of the data was assessed using the Shapiro-Wilk test. Non-normally distributed data was analyzed using the non-parametric Mann-Whitney test, while for the normal data was used the parametric Student’s t-test, one-way ANOVA followed by Dunnet multiple comparisons post-test and two-way ANOVA followed by Fisher’s LSD test were used in data analysis as detailed in the corresponding figure legend.

## Results

Previous research on cellular metabolic regulation in response to mechanical cues has primarily focused on short-term effects (24 hours). In this study, we evaluated the impact of long-term (72 hours) mechanical cues mimicking extracellular matrix (ECM) soft conditions (**Fig. 5.1A**) on mitochondrial metabolism, redox homeostasis and autophagy of neuronal cells.

### Soft ECM induces mitochondrial bioenergetic impairment

Cellular and mitochondrial metabolism has been increasingly recognized as a process influenced by mechanical signals [19, 20]. We start to investigate the generality of mitochondrial bioenergetic regulation in response to extracellular forces. This was achieved by measuring the real-time oxygen consumption rate (OCR) of HT22 cells (**Fig. 1B**). This is an *in vitro* model widely used for studying mechanisms of oxidative cell death associated with neurodegenerative diseases. Cellular OCR was consistently lower in a time-dependent manner in response to a soft ECM (2.5 kPa), with the largest difference in comparison to the stiffer ECM found after 72 hours (**Fig. 1C**). Although a decrease in cell proliferation has been described in response to extracellular forces [20, 34-37], HT22 cell density (**Fig. S1A**), nuclei number (**Fig. S1B**), and total protein content (**Fig. S1C**) did not show significant differences in response to 72 hours of incubation in soft ECM. Besides OCR, mitochondrial membrane potential (ΔΨm) and morphology can be indicators of mitochondrial (dys)homeostasis (**Fig. 1D**) [38]. HT22 cells presented a significant reduction in mitochondrial network area (**Fig. 1E**) and ΔΨm per cell (**Fig. 1F**), as demonstrated by the significant decrease in MitoTracker Green and MitoTracker Red fluorescence intensity, respectively. Additionally, we plotted the average data of mitochondrial network area against the average mitochondrial bioenergetic data (membrane polarization and oxygen consumption), which reveals a non-proportional decrease in both mitochondrial area, and membrane potential and OCR in HT22 cells in response to a soft ECM (**Fig. 1G**). Together, the data suggest a modulation in mitochondrial function (OCR and ΔΨm) in HT22 cells in response to long-term (72h) soft ECM mechanical cues.

**Figure 1.**
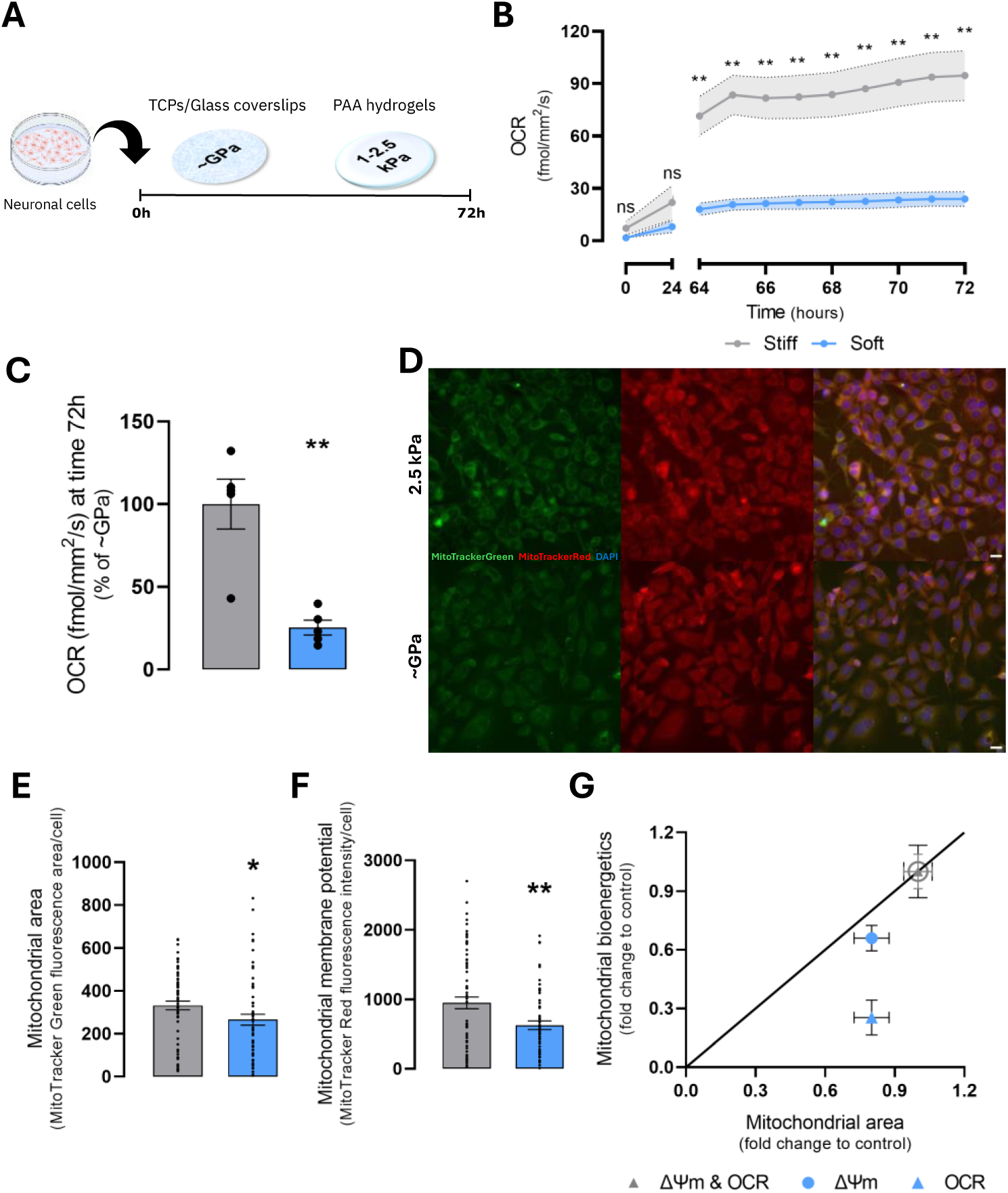
ECM mechanical cues regulate HT22 mitochondrial bioenergetics. **(A)** Neuronal cell lines were cultured on hydrogels resembling soft ECM (1-2.5 kPa) for 72 hours. **(B)** Real-time oxygen consumption rate (OCR) of HT22 cells cultured in soft ECM conditions was assessed through variations in using the Resipher. **(C)** Basal OCR of HT22 cells cultured during 72 hours in response to ECM mechanical cues. **(D)** Representative background-corrected images of HT22 cells stained with MitoTracker Green, MitoTracker Red and Hoechst (blue) to assess mitochondrial area and polarization in response to ECM mechanical cues. Scale bar: 20 μm. **(E)** Mitochondrial network area and **(F)** mitochondrial polarization (ΔΨm) in response to ECM mechanical cues were quantified through the MitoTracker Green and MitoTracker Red signal intensity per cell, respectively. **(G)** Average mitochondrial network area (MitoTracker Green) was plotted against the average mitochondrial bioenergetics (MitoTrackerRed and basal OCR), confirming a decrease in both mitochondrial area and bioenergetics (OCR and polarization, i.e.) in HT22 cells cultured on ECM soft substrate. Basal OCR traces represent the mean ± SEM, and the MitoTracker Green and Mitotracker Red stainings were analyzed from 10 to 14 fields per independent experiment in HT22 cells cultured on ECM soft substrate. **Statistics:** Data obtained from HT22 cells cultured on ECM soft substrate (2.5 kPa) (N=4) were compared to control conditions (∼GPa) (N=4) using a Student’s t-test to compare two mean values. Real-time OCR data was analyzed using two-way ANOVA, followed by Fisher’s LSD test for multiple comparisons. Significant differences between indicated conditions are marked by * (P<0.05) and ** (P<0.01).

### Soft ECM induces cellular redox (dys)homeostasis

Prompted by such widespread connection between mitochondria membrane polarization and oxygen consumption and extracellular forces, we explored the possibility that the alterations observed in mitochondrial function impact cellular energy status and redox state [39] in response to a soft ECM. Notably, soft ECM mechanical cues significantly reduced HT22 cellular metabolic activity (**Fig. 2A**) and ATP content (**Fig. 2B**). Moreover, the measurement of redox sensitive dye CM-H_2_DCFDA oxidation revealed a significant decrease in ROS levels in HT22 cells in response to a soft ECM (**Fig. 2C**). The nicotinamide adenine dinucleotide (NAD) and nicotinamide adenine dinucleotide phosphate (NADP) systems along with thiol/disulfide systems are central to redox control [40]. Interestingly, soft ECM mechanical cues significantly reduced HT22 cellular NAD^+^ and NADH (**Fig. 2D**), NADP and NADPH (**Fig. 2E**), and glutathione (GSH) (**Fig. 2F**) content. Together, these data suggest a cellular redox (dys)homeostasis prompted by lower antioxidant capacity, which may result in a more oxidative state of HT22 cells in response to long-term (72 hours) soft ECM mechanical cues.

**Figure 2.**
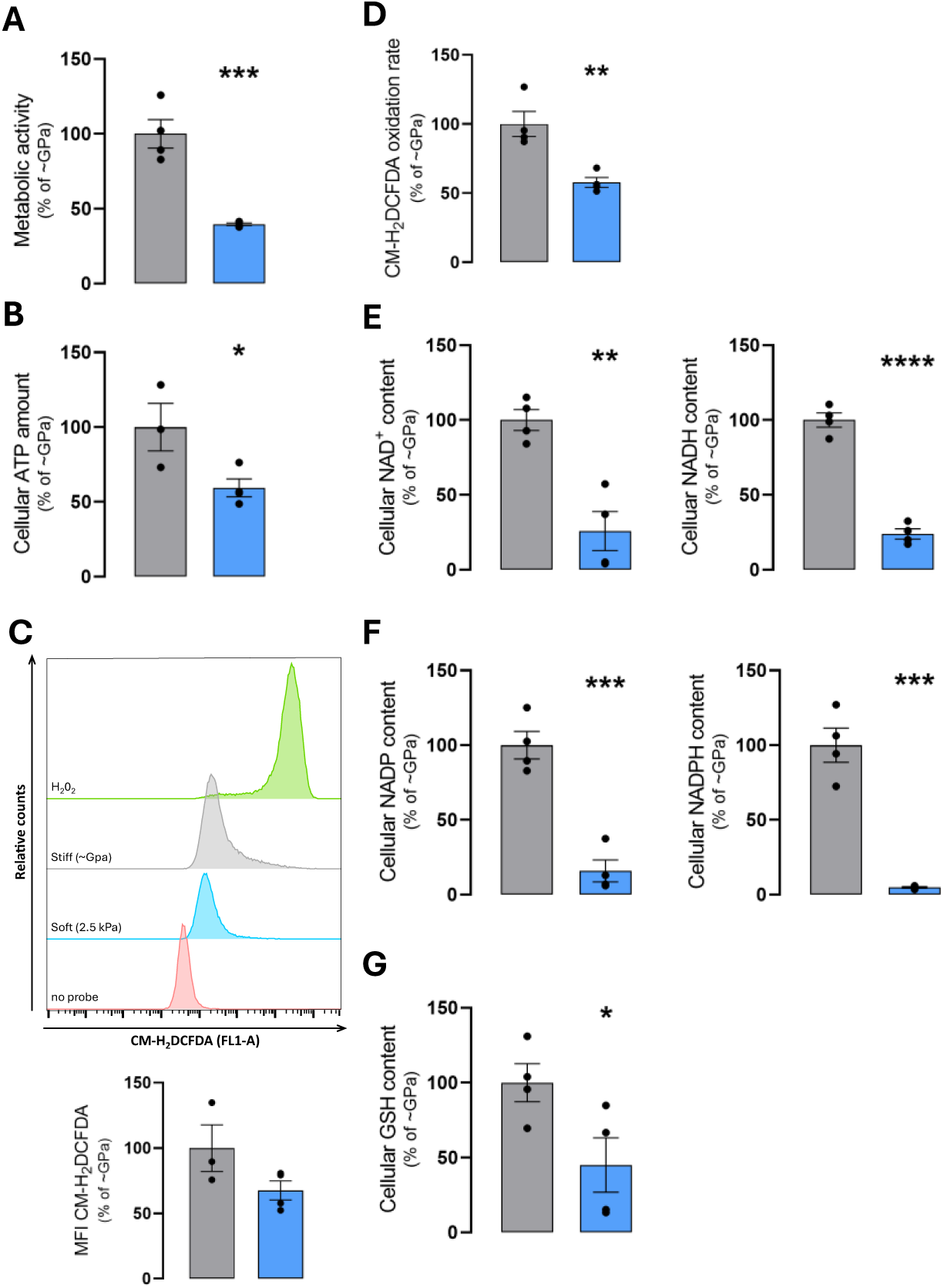
ECM mechanical cues regulate HT22 cellular redox status. **(A)** Cellular metabolic activity, **(B)** ATP content, **(C)** CM-H_2_DCFDA levels assessment by flow cytometry (oxidative stress), **(D)** NAD+, NADH, **(E)** NADP, NADPH, and **(F)** GSH content of HT22 cells cultured in soft ECM conditions (2.5 kPa) for 72 hours. **Statistics:** Data are presented as the mean ± SEM (N=4), and the results normalized to the control condition (∼GPa, i.e., stiff = 100%). Data obtained from HT22 cells cultured on ECM soft substrate (2.5 kPa) were compared to control conditions (∼GPa) using a Student’s *t*-test to compare two mean values. Significant differences between the indicated conditions are marked by * (*P*<0.05), ** (*P*<0.01), *** (*P*<0.001) and *** (*P*<0.0001).

### Soft ECM impacts neuronal auto(mito)phagy

The rising risk of cell death due to cellular oxidative stress and accumulation of impaired mitochondria can be prevented by the activation of autophagy / mitophagy signaling pathways [41, 42]. Thus, we next evaluated the impact of soft ECM on protein markers of cellular auto(mito)phagy in HT22 cells. The western blot analysis showed that soft ECM mechanical cues had no significant impact on autophagy-related proteins (**Fig. 3A, H)** P-MTOR and MTOR (**Fig. 3B**), beclin (**Fig. 3C**), SQSTM1 (**Fig. 3E**), LC3I (**Fig. 3F**), PINK1 (**Fig. 3I**), and Parkin (**Fig. 3J**) levels, but significantly increased ATG7 (**Fig. 3D**) and LC3 II (**Fig. 3G**) protein levels in HT22 cells. We next used two other types of neuronal-like cells to confirm whether the effects could also be observed.

**Figure 3.**
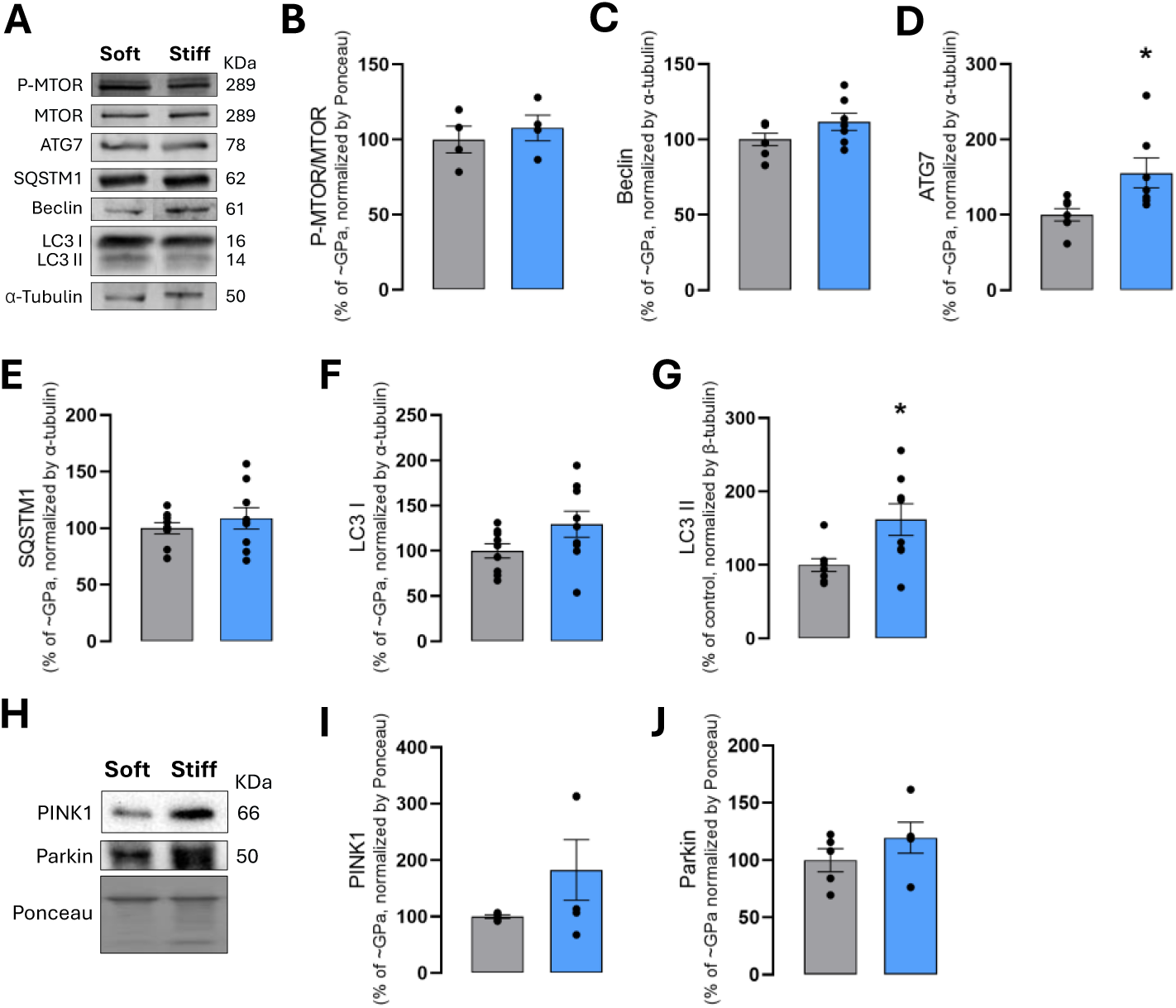
ECM mechanical cues impact on HT22 cellular auto(mito)phagy markers. (A, **H)** Representative Western blot result of whole-cell homogenates showing the cytosolic levels of proteins associated with autophagy **(B)** P-MTOR, MTOR, **(C)** Beclin, **(D)** ATG7, **(E)** SQSTM1, LC3 I, **(G)** LC3 II, and mitophagy pathways **(I)** PINK1 and **(J)** Parkin in HT22 cells cultured in soft ECM conditions (2.5 kPa) for 72 hours. These blots were contrast-optimized for visualization purposes. Quantification of the bands was performed using the original blots. Quantification of protein levels in multiple experiments (N=4-9) was normalized to α-tubulin levels or total amount of protein (Ponceau S staining), as indicated, and to the control condition (∼GPa, i.e., stiff = 100%). **Statistics:** Data obtained from HT22 cells cultured on ECM soft substrate (2.5 kPa) were compared to control conditions (∼GPa) using the Student’s *t*-test for comparison of two mean values. Significant differences between the indicated conditions are marked by * (*P*<0.05).

Protein markers of cellular auto(mito)phagy were also observed in human neuroblastoma SHSY-5Y cells in response to extracellular forces (1 kPa) (**Fig. S2A**). The western blot analysis showed that soft ECM mechanical cues had no significant impact on autophagy-related protein beclin (**Fig. S2B**), LC3I (**Fig. S2E**) and LC3II (**Fig. S2F**) levels, while significantly increased ATG7 (**Fig. S2C**) and decreased SQSTM1 (**Fig. S2D**) protein levels in SHSY-5Y cells. Primary mouse neurons were similarly cultured on hydrogels resembling soft ECM (2.5 kPa) for 6 days (**Fig. S3A, B**) ensuring the exclusion of glial cells from the culture, and protein markers of cellular auto(mito)phagy were also evaluated (**Fig. S3C**). The western blot analysis showed that autophagy-related protein beclin (**Fig. S3D**), ATG7 (**Fig. S3E**), SQSTM1 (**Fig. S3F**), LC3I (**Fig. S3G**), and LC3II (**Fig. S3H**) levels remained unchanged in primary mouse neurons in response to ECM mechanical cues. Altogether, data suggest that protein markers of autophagy catabolic processes are largely unaffected in response to long-term (72 h) soft ECM mechanical cues, an effect consistent in various neuronal cell lines tested.

### RhoA activation restores the YAP/TAZ mechanotransduction signaling pathway in neuronal cells

YAP inhibition in response to soft ECM and decreased actomyosin tension has been described in different cell types [19, 35, 36]. Thus, we next explored the possibility of restoring the YAP/TAZ mechanotransduction signaling pathway – typically associated with stiffer ECM – and its impact on autophagy, through the activation of RhoA, which recapitulates the increased cell contractility observed on stiff substrates. (**Fig. 4A**). The ratio of nuclear-to-cytoplasmic localization of mechano-responsive transcription factor YAP is significantly reduced in HT22 cells in response to soft ECM, an effect that was significantly rescued with the RhoA activator CN03 (1μg/mL, 3h) (**Fig. 4B**). Additionally, western blot analysis showed that ECM mechanical cues significantly decreased HT22 cells P-cofilin/Cofilin ratio, an indicator of mechanotransduction and actin cytoskeleton stabilization [43]. Conversely, RhoA activation with CN03 partially rescued the observed effects in HT22 cells (**Fig. 4C, D**). On the other hand, western blot analysis showed that RhoA activation with CN03 did not alter the protein content of autophagy-related proteins SQSTM1, LC3I, and LC3II (**Fig. 4C, E**) in HT22 cells in response to ECM mechanical cues. Similarly, RhoA activator CN03 significantly rescued the observed decrease in P-cofilin/Cofilin ratio, an indicator of mechanotransduction and actin cytoskeleton stabilization, in SHSY-5Y cells in response to soft ECM mechanical cues (**Fig. 4F, G**). However, RhoA activation by CN03 did not alter the protein content of autophagy-related proteins in SHSY-5Y cells in response to ECM mechanical cues. (**Fig. 4F, H**). These results suggest an inability of neuronal mechanosensitive cells to promote autophagic catabolic processes in response to long-term (72 hours) soft ECM mechanical cues.

**Figure 4.**
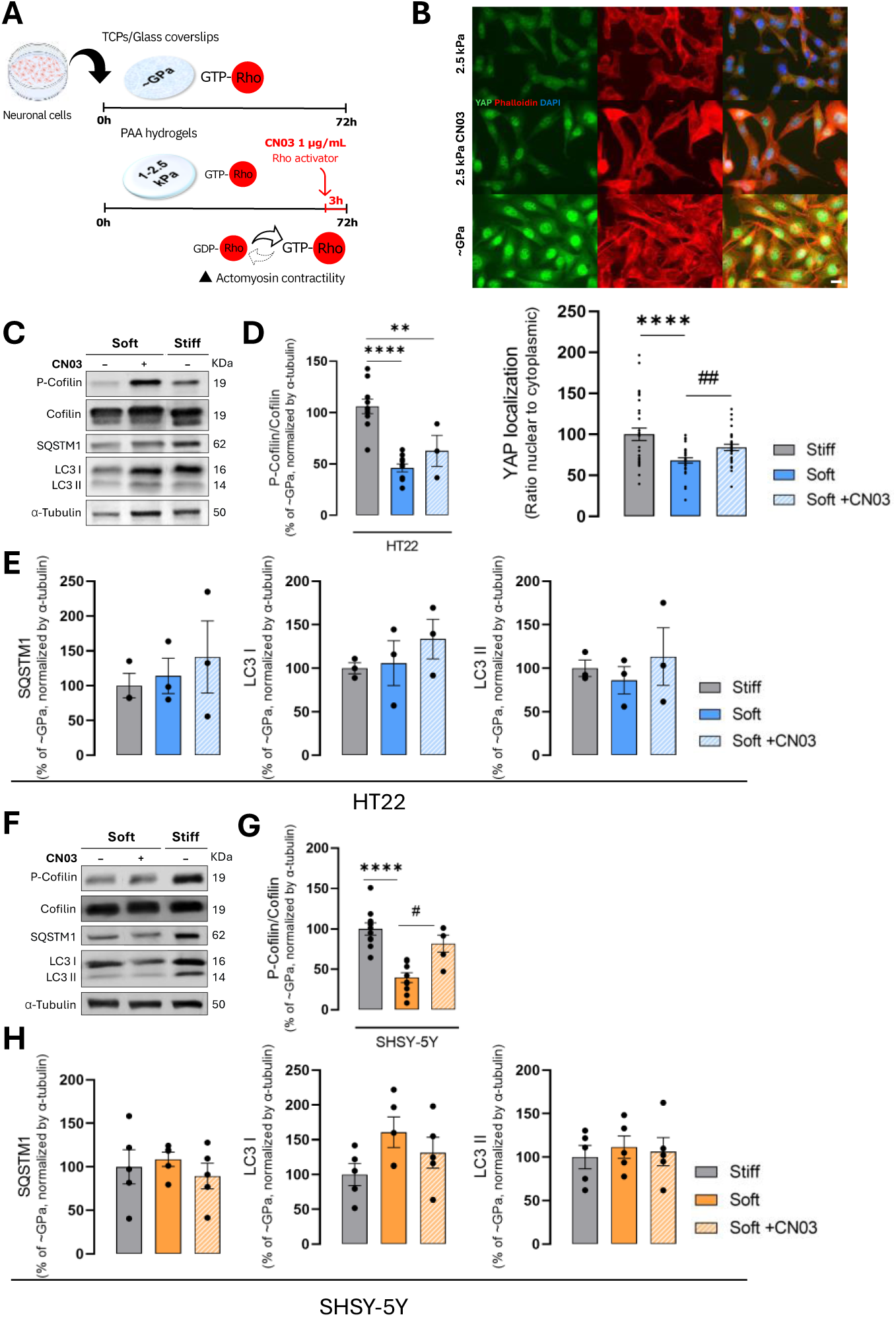
Restoration of mechanotransduction signaling pathways impact on cellular autophagy. **(A)** Mouse neuronal hippocampal HT22 cell line was cultured on hydrogels resembling soft ECM (2.5 kPa) for 72 hours in the presence or absence of RhoA activator CN03 (1 μg/mL, 3 hours) (Cytoskeleton, CN03), which recapitulates the increased cell contractility observed on a stiff substrate. **(B)** Representative images of cells stained with anti-YAP (green), Phalloidin (red) and DAPI (blue) and quantification of subcellular localization of YAP (ratio of nuclear-to-cytoplasmic) in HT22 cells cultured on soft ECM conditions (2.5 kPa) in the presence or absence of CN03. Scale bar: 20 μm. Quantification was performed from 10 to 12 fields per independent experiments HT22 cells cultured on ECM soft substrate. **(C)** Representative Western blot result of whole-cell homogenates showing the cytosolic levels of mechanotransduction-related **(D)** P-cofilin and cofilin, and **(E)** autophagy-related SQSTM1, LC3 I and LC3 II proteins in HT22 cells cultured on soft ECM conditions (2.5 kPa) in the presence or absence of CN03. **(F)** Representative Western blot result of whole-cell homogenates showing the cytosolic levels of mechanotransduction-related **(G)** P-cofilin and cofilin, and **(H)** autophagy-related SQSTM1, LC3 I and LC3 II proteins in SHSY-5Y cells cultured on soft ECM conditions (1 kPa) in the presence or absence of CN03. The blots were contrast-optimized for visualization purposes. Quantification of the bands was performed using the original blots (N=3-10). Quantification of protein levels in multiple experiments was normalized to α-tubulin levels and to the control condition (∼GPa, i.e., stiff = 100%). **Statistics:** Data obtained from HT22 or SHSY-5Y cells cultured on Soft (2.5 kPa or 1 kPa, respectively) +CN03 was compared to Soft using the Student’s *t*-test for comparison of two mean values, and the one-way ANOVA with Dunnett multiple comparison post-test to compare more than two groups with one independent variable. Significant differences between the indicated conditions are marked by # (*P*<0.05), ## (*P*<0.01), ** (*P*<0.01) and ****(*P*<0.0001).

### Soft ECM disrupts neuronal cells’ autophagic flux

We next sought to better understand the nature and extent of the alterations identified on neuronal cells catabolic processes on soft ECM, focusing on autophagic flux (**Fig. 5A**). We first focused on evaluating LC3 II levels, in the absence and presence of chloroquine (ChQ 100 μM, 3 h) an inhibitor of autolysosomal degradation [44, 45]. LC3 II is degraded during the final stage of autophagy, but with chloroquine-induced blockade of autolysosomal degradation, LC3 II accumulates within cells [44]. Chloroquine induced a significant increase of LC3 II protein levels in HT22 cells in response to extracellular forces to a similar extent when compared to control conditions (∼GPa) (**Fig. 5B**). However, LC3II turnover is significantly decreased in response to soft ECM mechanical cues (**Fig. 5C**), indicating an impairment in HT22 maximal autophagic capacity. Similarly, chloroquine induced a significant increase of LC3 II protein levels in SHSY-5Y cells on soft conditions, which is significantly different when compared to control conditions (∼GPa +ChQ) (**Fig. 5D**). However, LC3II turnover remained unchanged in SHSY-5Y cells in response to extracellular forces (**Fig. 5E**). Interestingly, the same pattern was observed on LC3 II protein levels (**Fig. 5F**) and LC3II turnover (**Fig. 5G**) of mouse primary neurons in response to extracellular forces. Collectively, these findings suggest long-term (72 hours) soft ECM mechanical cues impairs maximal autophagic capacity of neuronal mechanosensitive cells, which probably reflect an (im)balance between autophagosome formation and degradation, the leading cause for impairment of mitochondrial function and cellular redox (dys)homeostasis.

**Figure 5.**
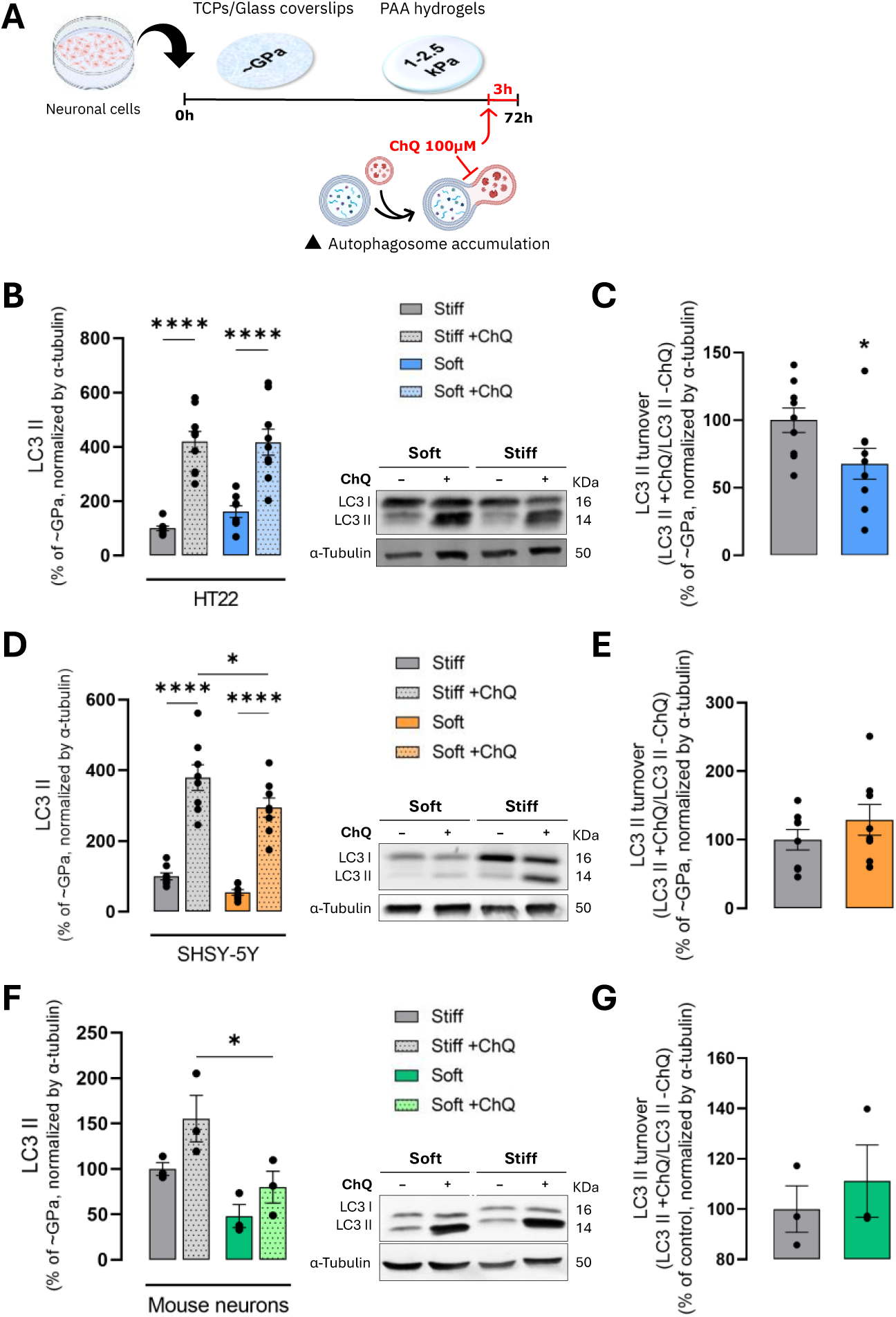
ECM mechanical cues disrupt neuronal cells’ autophagic flux. **(A)** Mouse neuronal hippocampal HT22 cells and primary neurons and human neuroblastoma SHSY-5Y cells were cultured on hydrogels resembling soft ECM (2.5 kPa and 1 kPa, respectively) for 72 hours in the presence or absence of chloroquine (ChQ; 100 μM, 3 hours), a lysosomal protein degradation inhibitor. **(B)** Representative Western blot result of whole-cell homogenates showing the cytosolic levels of LC3 II protein in HT22 cells cultured on soft (2.5 kPa) and stiff (∼GPa) ECM conditions in the presence or absence of ChQ. **(C)** LC3 II turnover in HT22 cells cultured on soft ECM conditions (2.5 kPa). **(D)** Representative Western blot result of whole-cell homogenates showing the cytosolic levels of LC3 II protein in SHSY-5Y cells cultured on soft (1 kPa) and stiff (∼GPa) ECM conditions in the presence or absence of ChQ. **(E)** LC3 II turnover in SHSY-5Y cells cultured on soft ECM conditions (1 kPa). **(F)** Representative Western blot result of whole-cell homogenates showing the cytosolic levels of LC3 II protein in mouse primary neurons cultured on soft (2.5 kPa) and stiff (∼GPa) ECM conditions in the presence or absence of ChQ. **(G)** LC3 II turnover in mouse primary neurons cultured on soft ECM conditions (2.5 kPa). The blots were contrast-optimized for visualization purposes. Quantification of the bands was performed using the original blots (N=3-9). Quantification of protein levels in multiple experiments was normalized to α-tubulin levels and control condition (∼GPa, i.e., stiff = 100%). **Statistics:** Data obtained from mouse HT22 and primary neurons or SHSY-5Y cells cultured on ECM soft substrate (2.5 kPa or 1 kPa, respectively) in the presence of ChQ were compared to control conditions (∼GPa) using two-way ANOVA with Tukey multiple comparison post-test, and data obtained from LC3 II turnover was compared using the Student’s *t*-test for comparison of two mean values. Significant differences between the indicated conditions are marked by * (P<0.05) and ****(P<0.0001).

## Discussion

The specific composition of extracellular matrix (ECM) of different tissues and organs is unique, giving rise to their characteristic biomechanical properties. However, studies incorporating ECM mechanics in the neuronal context are primarily focused on enhancing neuronal differentiation and growth to support treatments for brain injuries and neurodegenerative disorders [5, 6, 8, 46, 47]. Although soft ECM can enhance hormetic and neuroprotective effects of low-dose topoisomerase inhibitors in neuronal PC12 cells, potentially through the EGFR/PI3K/AKT signaling pathway [48], studies on cellular metabolic regulation in response to mechanical cues have primarily focused on evaluate the short-term (24 hours) impact of ECM stiffness in cancer cells [4, 18-20, 23, 25, 26]. In this study, we directly address how neuronal cells’ metabolic processes respond to low actomyosin tension favoring conditions (soft ECM), expanding the scope of current research on ECM-mediated signaling in neuronal cells. In light of this, our study aimed to provide new insights into the effects of longer-term (72 hours) exposure to soft ECM mechanical cues on neuronal cell homeostasis, including mitochondrial membrane potential and oxygen consumption, redox homeostasis, and autophagy.

Our findings reveal that longer-term exposure to a soft ECM compromises mitochondrial function, cellular metabolic activity and energy production of neuronal cells, which is accompanied by a redox imbalance. Notably, no significant differences in nuclear number or total protein content were observed in HT22 neuronal cell line in response to decreased extracellular forces, suggesting that longer-term ECM softening is unlikely to impact cell proliferation. Several evidence have been reporting the cell-type dependent impact of ECM stiffness on cell proliferation. Soft ECM decrease proliferation of fibroblasts, epithelial cells, and various cancer cells [20, 34-37], while increased proliferative rates under softer ECM conditions have been described in myoblasts, spinal cord progenitor cells and neural progenitor cells [47, 49-51]. This suggests that the relationship between ECM stiffness and cell proliferation is not universally linear, probably due to differences in cell origin and/or energy requirements, but it is unlikely to affect our main findings. On the other hand, cells with reduced actomyosin tension have been reported to have decreased mitochondrial energy production activity [20, 21], consequently with lower levels of ATP/ADP ratio compared with higher intracellular tension [20, 21]. These results align with observations in cancer cells experiencing short-term (24 h) low adhesion and traction forces [21] or exposed to soft ECM [20], supporting the notion that reduced actomyosin tension negatively impacts mitochondrial function. In contrast to neuronal cells, cancer cell lines exposed to soft ECM or reduced actomyosin tension have shown an increase in ROS production [20]. Notwithstanding, reducing the overall antioxidant capacity, namely decreased glutathione levels and nicotinamide adenine dinucleotides, is likely to promote a state of oxidative stress within these cells. Although the final outcome on soft ECM is an increased cellular oxidative status, our findings suggest that redox responses to ECM stiffness may slightly vary between cell types and are influenced by exposure duration. Neuronal cells, in particular, display a time-dependent sensitivity to ECM mechanical cues, underscoring their unique metabolic adaptations to ECM stiffness [5, 6].

The prolonged soft ECM conditions impact mitochondrial function and redox balance on neuronal cells, which may activate signaling pathways leading to cell death. Autophagy and mitophagy play a critical role in determining cellular outcomes in response to shifts in metabolic status, stress, and other intra- and extracellular cues [41, 42]. This catabolic process was described to also be activated in response to conditions that favor actomyosin tension, including ECM stiffness, via YAP/TAZ signaling pathway [25, 26, 52-54]. Notably, autophagy-related protein markers remained unchanged across different neuronal cell types following long-term exposure to soft ECM mechanical cues. Previous studies in cancer cell lines, epithelial cells and fibroblasts have reported short-term exposure to soft ECM impairs autophagosome formation, largely due to YAP/TAZ inhibition [25, 26]. In our study, however, pharmacological activation of the YAP/TAZ pathway in soft ECM did not show significant changes in autophagy-related protein markers. This finding suggests that neuronal mechanosensitive cells exposed to long-term soft ECM mechanical cues exhibit a fundamentally limited ability to regulate autophagy in response to these mechanical conditions.

Moreover, our data points to a reduced maximal autophagic capacity in response to long-term soft ECM in both mouse and human neuronal cells. This effect may arise from an imbalance in autophagosome formation and degradation, as shown by decreased LC3 II levels in HT22, SH-SY5Y cells, and primary mouse neurons following long-term soft ECM exposure, which became evident only when autophagic flux was inhibited with chloroquine. As a consequence of chloroquine-induced autolysosomal degradation inhibition, LC3 II levels in cells on soft ECM did not show the same degree of accumulation as those on stiff ECM, suggesting that long-term soft ECM exposure may impair autophagosome formation, thereby limiting the autophagic response in these cells. The observed autophagic imbalance may contribute to the impairment in mitochondrial function and the dysregulation of cellular redox homeostasis, suggesting that autophagic insufficiency could be a key factor linking mechanical cues to mitochondrial and oxidative stress-related dysfunction in neuronal cells.

This study provides new evidence that mechanical properties of the ECM play a critical role in the metabolic homeostasis of neuronal cells, specifically impacting autophagic capacity, mitochondrial and redox regulation pathways in a time-dependent manner (**Fig. 6**). We validated the longer-term soft ECM exposure effects across both human and mouse neuronal cells, demonstrating that the observed cellular responses are broadly applicable across species. Furthermore, given that different cell lines exhibit unique metabolic and signaling profiles, assessing neuroblastoma cells (SH-SY5Y), a hippocampal cell line (HT22), and primary mouse neurons highlights the relevance of these findings for neurodegenerative disease models, enhancing their physiological significance. Our results emphasize the importance of incorporating mechanical cues into *in vitro* experiments to more accurately mimic *in vivo* environments, particularly when studying neurodegenerative diseases.

**Figure 6.**
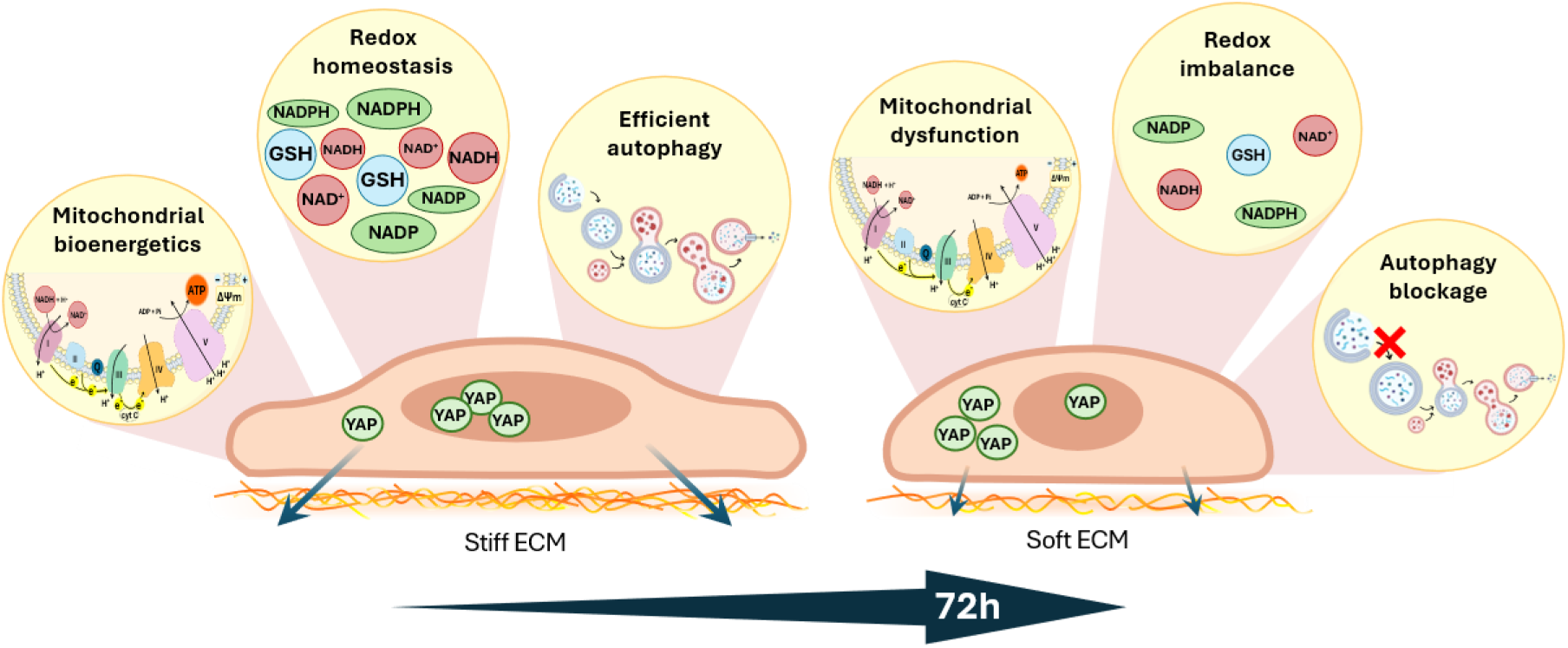
Neuronal cell’s metabolic response to low actomyosin tension favoring conditions (soft ECM). Longer-term (72 h) exposure to soft ECM resulted in mitochondrial bioenergetic dysfunction, redox imbalance, and autophagy blockage. Cells exposed to long-term soft ECM displayed reduced maximal autophagic capacity, likely due to disrupted autophagosome formation and degradation, as evidenced by decreased LC3 II levels following chloroquine-induced inhibition of autophagic flux. This autophagy impairment, combined with increased cellular oxidative stress, highlights the presence of metabolic alterations depending on ECM stiffness. **Abbreviations:** NAD(H), nicotinamide adenine dinucleotide; NADP(H), nicotinamide adenine dinucleotide phosphate; GSH, reduced glutathione; ATP, adenosine triphosphate; ADP, adenosine diphosphate; Pi, inorganic phosphate; cyt C, cytochrome C; YAP, Yes-associated protein; ECM, extracellular matrix.

## Conclusions

Developing a cellular model that faithfully mimics the pathology of neurodegenerative disorders has been a persistent challenge. By demonstrating that ECM stiffness directly affects metabolic processes in neuronal cells, our results highlight the limitations of standard *in vitro* models that do not account for ECM mechanics, which may lead to findings with reduced translational relevance. Therefore, integrating realistic mechanical conditions is essential for obtaining more accurate insights into the pathophysiology of neurodegenerative disorders by facilitating the precise replication of the cellular, biochemical, and physical characteristics of native tissue microenvironments [11, 14-16]. This study can provide new avenues to explore the complex intersections of mechanics and metabolism, enhancing the reliability of therapeutic assessments.

## CRediT authorship contribution statement

HG, Conceptualization, data curation, writing, editing and visualization; TL, data curation and methodology; JT, data curation; MF, review and editing; CA, Conceptualization and supervision; SS, data curation and methodology; LF, review and editing; CC, review and editing; PJO, supervision, review and editing; JT, Conceptualization, data curation, supervision, writing, review, editing and visualization; MG, Conceptualization, data curation, supervision, review and editing. All authors have read and agreed to the published version of the manuscript

## Data available within the article or its supplementary materials

The authors confirm that the data supporting the findings of this study are available within the article.

## Acknowledgements

This work was financed by the European Regional Development Fund (ERDF), through the Centro 2020 Regional Operational Programme and through the COMPETE 2020 - Operational Programme for Competitiveness and Internationalisation and Portuguese national funds via FCT, under project[s]: EXPL/BIA-BQM/1361/2021, UIDP/04539/2020 and LA/P/0058/2020. PAS GRAS project has received funding from the European Union’s Horizon Europe under grant agreement No 101080329. H. Gerardo (SFRH/BD/147316/2019 and COVID/BD/153559/2024) and J. Teixeira (2020.01560.CEECIND) acknowledge FCT, I.P. for the research contract.

