## Supplementary data for "Extracellular matrix mechanical cues (dys)regulate metabolic redox homeostasis due to impaired autophagic flux"

**Supplementary Figures**

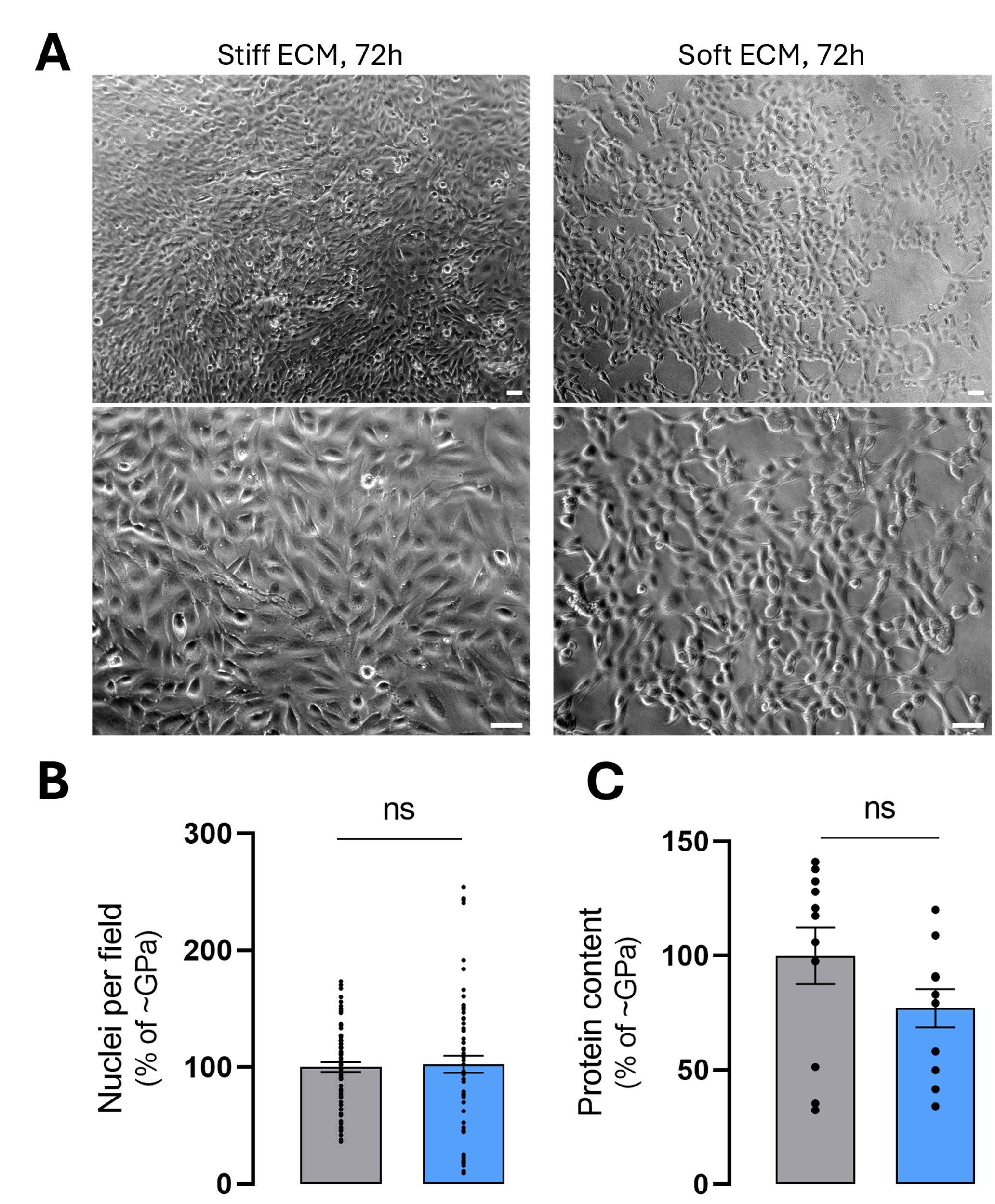

**Figure S1. ECM mechanical cues do not impact on HT22 cellular growth. (A)** Representative contrast phase images of mouse neuronal hippocampal HT22 cells after 72 hours in culture on stiff (~GPa) and soft (2.5 kPa) ECM, scale bar: 50 μm. **(B)** Nuclei number per filed (16 fields with 0.11 mm² per independent experiment, N=4) and **(C)** protein content quantification (N=11) from HT22 cells in same conditions. **Statistics:** Data obtained from HT22 cells cultured on ECM soft substrate (2.5 kPa) were compared to control conditions (~GPa) using the Mann-Witney test and Student’s *t*-test for comparison of two mean values.

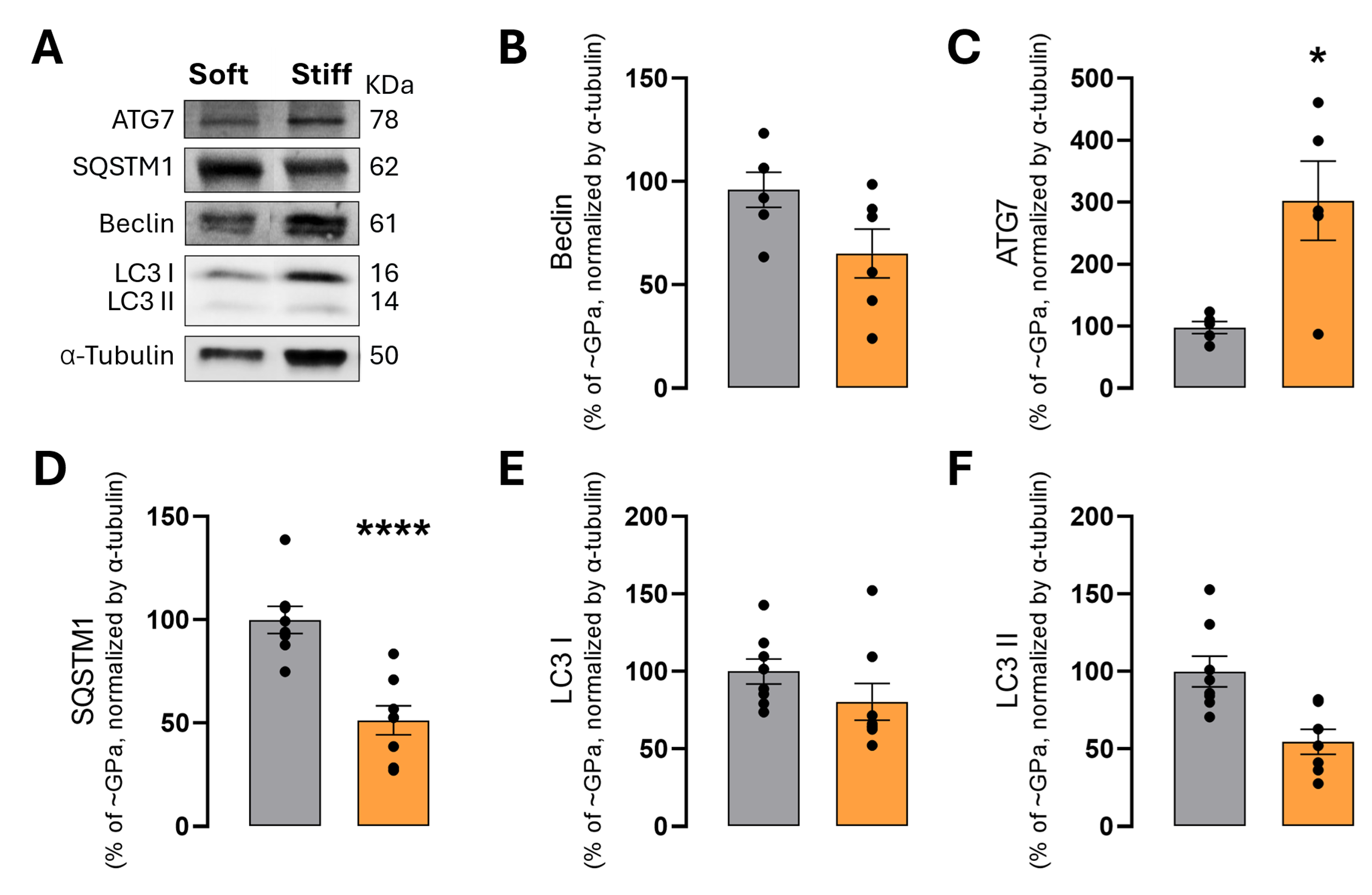

**Figure S2. ECM mechanical cues impact on SHSY-5Y cellular auto(mito)phagy markers. (A)** Representative Western blot result of whole-cell homogenates showing the cytosolic levels of proteins associated with autophagy **(B)** Beclin, **(C)** ATG7, **(D)** SQSTM1, **(E)** LC3 I, and **(F)** LC3 II in HT22 cells cultured in soft ECM conditions for 72 hours. These blots were contrast-optimized for visualization purposes. Quantification of the bands was performed using the original blots. Quantification of protein levels in multiple experiments (N=5-8) was normalized to α-tubulin levels or total amount of protein (Ponceau S staining), as indicated, and to the control condition (~GPa, i.e., stiff = 100%). **Statistics:** Data obtained from SHSY-5Y cells cultured on ECM soft substrate (1 kPa) were compared to control conditions (~GPa) using the Student’s *t*-test for comparison of two mean values. Significant differences between the indicated conditions are indicated by * (P<0.05) and **** (*P*<0.0001).

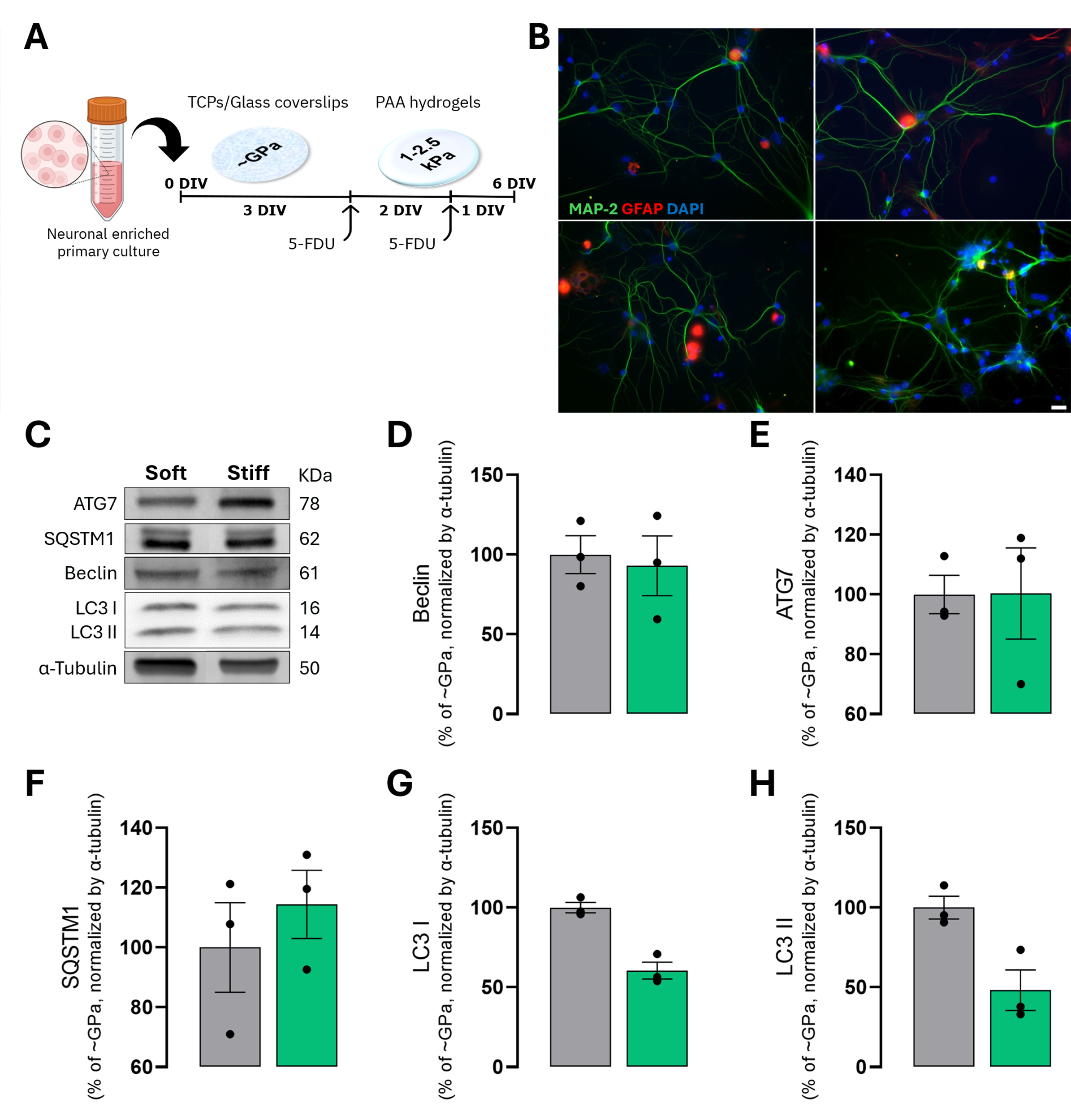

**Figure S3. ECM mechanical cues impact on mouse primary neurons auto(mito)phagy markers. (A)** Mouse neuronal enriched primary cell suspension was directly plated on hydrogels resembling soft ECM (2.5 kPa) and stiff ECM (~GPa) and cultured for 6 days in vitro (DIV). **(B)** Representative background-corrected images of primary mouse neurons cultured on hydrogels resembling soft ECM (2.5 kPa) for 6 days. Cells stained with anti-MAP-2 (green) to identify neurons, GFAP (red) to identify astrocytes and DAPI (blue) for nuclear staining. **(C)** Representative Western blot result of whole-cell homogenates showing the cytosolic levels of proteins associated with autophagy **(D)** Beclin, **(E)** ATG7, **(F)** SQSTM1, **(G)** LC3 I, and **(H)** LC3 II in mouse primary neurons cultured in soft (2.5 kPa) and stiff (~GPa) ECM conditions for 6 days. These blots were contrast-optimized for visualization purposes. Quantification of the bands was performed using the original blots. Quantification of protein levels in multiple experiments (N=3) was normalized to α-tubulin levels and to the control condition (~GPa, i.e., stiff = 100%). **Statistics:** Data obtained from mouse primary neurons cells cultured on ECM soft substrate (2.5 kPa) were compared to control conditions (~GPa) using the Student’s *t*-test for comparison of two mean values.
